# A novel pollen-tracking method: using quantum dots as pollen labels

**DOI:** 10.1101/286047

**Authors:** Corneile Minnaar, Bruce Anderson

## Abstract

To understand the evolution of flowers and mating systems in animal-pollinated plants, we have to directly address the function for which flowers evolved—the movement of pollen from anthers to stigmas. However, despite a long history of making significant advances in our understanding natural selection and evolution, the field of pollination biology has largely studied pollen movement indirectly (e.g., pollen analogues or paternity assignment to seeds) due to a lack of suitable pollen tracking methods. Here, we develop and test a novel pollen-tracking technique using quantum dots as pollen-grain labels. Quantum dots are semiconductor nanocrystals so small in size that they behave like artificial atoms. When exposed to UV light, they emit extremely bright light in a range of different colours. Their photostability, broad excitation range, and customisable binding-ligands make quantum dots ideal bio-labels. We tested the suitability of CuInSe_x_S_2-x_/ZnS (core/shell) quantum dots with oleic acid (zinc-oleate complex) ligands as pollen-grain labels. We found that quantum dots attach to pollen grains of four different species even after agitation in a polar solvent, suggesting that the oleic acid ligands on quantum dots bind to pollenkitt surrounding pollen grains. We also showed that most pollen grains within anthers of the same four species are labelled with quantum dots after applying sufficient quantum-dot solution to anthers. To test whether quantum-dot pollen-labels influenced pollen transport, we conducted pollen transfer trials on *Sparaxis villosa* (Iridaceae) using captively reared honeybees. We found no difference in pollen transport between labelled and unlabelled pollen grains. Our experiments therefore demonstrate the potential for quantum dots to be used as easily applied pollen labels, which allow subsequent tracking of the fates of pollen grains in the field. The ability to track pollen grain movement in situ, may finally allow us to address an historically neglected aspect of plant reproductive ecology and evolution.

## INTRODUCTION

Floral biology has contributed significantly to our understanding of evolution for more than three centuries through studies of trait inheritance (Mendel, 1866), sexual reproduction (e.g., Grew 1682; Camerarius 1694), the effects of inbreeding and outcrossing (e.g., Knight 1799; Darwin 1876; Levin 1984), mating systems (e.g., Darwin 1877; Barrett 1977), local adaptation (e.g., Robertson and Wyatt 1990; Newman *et al*. 2015), and coevolution (e.g., Nilsson 1988; Alexandersson and Johnson 2002; Anderson and Johnson 2008; Pauw *et al*. 2009). Remarkably, many of these studies, spanning 333 years, used the same basic techniques, highlighting the suitability of flowering plants as subjects in low-tech evolutionary experiments: their seeds are easy to store, transport, and grow; controlled crosses can easily be achieved through hand pollination; variations in floral and pollinator traits are easy to quantify; floral interactions with pollinators are easy to observe; and perhaps most important to evolutionary biologists, reproductive fitness can be quantified by simply counting the number of seeds produced by a flower. However, this low-tech approach has left the field somewhat stagnant in one very important aspect of evolutionary pollination biology—the quantification of pollen movement and male fitness.

Flowers have evolved for no other purpose than to serve the basic requirements for male and female reproductive success—pollen export and pollen import, respectively. Without the movement of pollen from anthers to stigmas, no reproduction would occur. Therefore, the study of floral evolution should be grounded on studies of pollen movement. Yet, despite over a century of fundamental work on floral evolution, our technical inability to track pollen grains has rendered direct assessments of pollen movement one of the most poorly studied aspects of plant reproductive biology. While the relatively recent development of paternity analyses can tell us which pollen grains sired seeds (Jones *et al*., 2010), we still do not know how these successful pollen grains arrived on stigmas to begin with. Consequently, nearly all studies of plant mating have been limited to post-fertilisation measures of mating success, making the methods of studying plant reproduction analogous to studying animal reproduction without accounting for the mechanistic links between mating behaviour and reproductive success. On average, less than 2% of pollen grains produced by flowers successfully reach conspecific stigmas (Harder and Thomson, 1989; Gong and Huang, 2014). The remaining 98% of grains are lost to multiple theoretical fates (Inouye *et al*., 1994), with each avenue of pollen loss representing an opportunity for selection to act on different floral traits that reduce the amount of wasted reproductive potential (Harder and Thomson, 1989; Harder and Barrett, 1996). The transport of pollen from anthers to stigmas is clearly an important phase for selection on floral traits, and therefore floral evolution. To fully understand the evolutionary function of floral traits, we need to be able to track pollen grains—not just those that successfully fertilise ovules—but all pollen grains.

### Pollen tracking methods to date

Perhaps the best success in tracking pollen comes from a single plant family, the Orchidaceae. Orchid pollen grains are contained in large pollen packets called pollinaria, which can be stained using dyes (Peakall, 1989) or labelled using uniquely coded microfilm tags (Nilsson *et al*., 1992). The dyed/labelled massulae (subunits of a pollinarium) can then be recovered and counted from other flowers once transferred (Peakall, 1989; Nilsson *et al*., 1992; Johnson *et al*., 2005; Jersáková and Johnson, 2007). However, most angiosperms produce granular pollen, and attempts at staining entire anthers containing granular pollen grains with dye have largely failed as pollen grains do not absorb dyes well in situ. This method of pollen staining has therefore rarely been used [only twice to our knowledge: Huang and Shi (2013); Armbruster *et al*. (2014) since the first attempt (Huang and Guo, 1999)]. Pollen grains have also been labelled with radioactive elements (Colwell, 1951), neutron-activated elements (Gaudreau and Hardin, 1974; Handel, 1976), and ^14^C labels (Reinke and Bloom, 1979; Pleasants *et al*., 1990). However, concerns about environmental exposure to radioactive labels, the complicated and time-consuming process of detection of neutron-activated and ^14^C labels (up to 14 weeks), and the limited number of one unique labels (usually just one), ultimately rendered these methods ineffective.

Instead of attempting to directly label pollen grains, some researchers have used fluorescent dye powder as a pollen proxy (Stockhouse II, 1976; Price and Waser, 1982; Waser and Price, 1982). In some cases, fluorescent dye particle deposition on stigmas correlates relatively well with pollen grains deposited per visit (Waser and Price, 1982; Fenster *et al*., 1996; Van Rossum *et al*., 2011). However, pollen grains are often found on stigmas when dye is not, and vice versa (Waser and Price, 1982). Studies have also found that dye particles can significantly over- or underestimate pollen transfer for different pollinators species (Thomson *et al*., 1986; Waser, 1988; Campbell, 1991; Adler and Irwin, 2006). Therefore, fluorescent dye particles cannot be used to estimate numbers of pollen grains transferred without first performing labour-intensive dye–pollen transfer comparisons (using the species of interest in the study) to determine the relationship between dye particle counts and pollen grain counts. Such experiments require flight cages and captive, pollen-free pollinators, which is impractical for most study systems. A similar method used micronized metal (Zn and Sn) dusts applied to dehisced anthers (Wolfe *et al*. 1991). While some of the metal particles labelled grains directly, their presence on pollen grains was likely superficial. Moreover, to detect metal dust particles, samples have to be gold-plated for subsequent scanning electron microscopy. To our knowledge, this method has never been applied outside of the original study which reported it.

Other researchers have effectively managed to use intraspecific colour variations of pollen grains to track pollen in a few limited cases where these variations exist (e.g., Thomson and Plowright 1980; Holsinger and Thomson 1994). Similarly, intraspecific variation in pollen size associated with upper and lower anthers in distylous morphs have been exploited to quantify pollen movement (Nichols, 1985; Stone, 1995). Unfortunately, these methods are limited in their applicability as most other systems do not have pollen colour or size polymorphisms which can be exploited. Recently, researchers have started to use the genetic labels already present in pollen grains. By microsatellite genotyping individual pollen grains (Matsuki *et al*., 2007) researchers can identify the individual plant origin of pollen grains found on floral visitors (Matsuki *et al*., 2008; Hasegawa *et al*., 2009, 2015) and stigmas (Hasegawa *et al*., 2009). However, this technique is incredibly labour intensive and expensive, requiring careful pollen isolation, DNA extraction, and sequencing of individual pollen grains. In studies requiring quantitative assessments of various aspects of the pollen export process, these techniques may not be. To make tracking pollen a practical and accessible to pollination biologists, we need to develop an easy, cost-effective method which can be applied in situ.

### Quantum dots as potential pollen labels

Quantum dots are extremely small nanoparticles made from semiconductor metals. They range in diameter from 2 to 10 nm (10 to 50 atoms across) (Neeleshwar *et al*., 2005) making them smaller than the smallest viruses (typically 20 nm in diameter) (Dimmock *et al*., 2016). Their composition and extremely small size results in tight confinement of electrons, or electron-holes, in bound discrete states similar to the behaviour of electrons and electron-holes in individual atoms (Gammon, 2000). This atom-like behaviour causes quantum dots to emit bright light in the visible spectrum when excited with UV radiation (Ekimov and Onushchenko, 1981, 1982; Brus, 1984). To understand how electrons behave in quantum dots, it is important to first understand how they behave in relatively large semiconductor metal objects: when excited (e.g., exposed to electricity) electrons bound to atoms (in the valence band), become free (jump to the conduction band) and can move within the crystal lattice of a large semiconductor object (i.e., electrical conduction) (Cho, 1979; Dean and Herbert, 1979). Electrons behave in this way as long as the semiconductor object remains large relative to the wavelength of the electrons (Brus, 1984). When the semiconductor metal object is shrunk to the nanoscale (quantum dots), the valence and conduction energy bands that the electrons can occupy become discrete (Yoffe, 1993, 2001). In a spherical quantum dot, this can be theoretically visualised as a ball with discrete layers: the outer layer is the conductance band, and the inner layer is the valence band. When electrons become excited (usually through UV radiation), they jump from the valence band to the conduction band (Ekimov, 1991), leaving behind an electron-hole (a quasiparticle with a positive charge relative to electrons) (Yoffe, 1993, 2001). Normally, in a large semiconductor object, the electron can move freely, independent of the electron-hole, but because of the tight confinement of the electron inside the quantum dot, the electron and the electron-hole are bound in close proximity and form an exciton which jumps to the conduction band (Cho, 1979; Dean and Herbert, 1979; Kusrayev, 2008). When the exciton returns to the ground state (valence band), it emits light energy, causing the quantum dot to fluoresce (Ekimov, 1991). The size of the quantum dot determines the radius of the two energy bands, and therefore the exciton’s light emission wavelength (Yoffe, 2001). The emission colour of quantum dots can therefore be tuned precisely by altering the size of the quantum dot.

Quantum dots were first employed as bio-labels two decades ago (Bruchez Jr., 1998; Chan, 1998) and offer several advantages over traditional bio-labels (Jaiswal and Simon, 2004): (1) Their colours are highly tuneable. (2) They have much greater photostability than traditional fluorescent markers; traditional fluorescent markers lose their fluorescence comparatively quickly under excitation. (3) They have very large Stokes-shifts (difference between excitation and emission wavelengths), and therefore multiple different coloured quantum dots can be excited by a single light source and detected simultaneously. In contrast, traditional fluorescent markers have small Stokes-shifts resulting in overlap between excitation and emission wavelengths for different coloured fluorophores. Therefore, different coloured fluorophores cannot be viewed simultaneously and require a unique set of emission and excitation filters for visualisation. (4) The ease with which bio-functional groups can be attached to quantum dots allows them to be used as bio-la-bels for virtually any biomolecule, including pollen grains.

Initially, quantum dots were made from toxic heavy-metal semiconductor cores (primarily Cadmium), which precluded their use in natural environments (Hardman, 2006). However, in recent years, several non-toxic alternatives have been developed (Xu *et al*., 2016), and many different forms are available commercially, making the use of quantum dots as pollen labels a practical possibility. Most applications of quantum dots as bio-labels attach highly specific functional groups to quantum dots to enable targeted labelling of molecules of interest. For quantum dots to work as universal pollen labels, a more general binding strategy is required. Many commercially available quantum dots carry oleic acid ligands which allow them to be dissolved in non-polar solvents. Oleic acid is lipophilic and may therefore bind to lipid-rich pollenkitt (Pacini and Hesse, 2005) surrounding pollen grains. Here, we evaluate the potential of quantum dots as pollen labels through three important criteria required for quantum dots to function as pollen-grain labels: (1) quantum dots must attach directly to pollen grains; (2) the application of quantum dots to an anther should result in most pollen grains being labelled; (3) quantum-dot labels should not affect pollen grain transport. In addition, to visualise or “read” quantum-dot pollen-labels, we designed and built a quantum-dot excitation box which converts any standard dissection microscope into a fluorescence microscope capable of detecting quantum-dot-labelled pollen grains on insects and stigmas.

## PROPOSED METHOD

### Quantum dots

For all experiments, we used heavy-metal-free CuInSe_x_S_2-x_/ZnS (core/shell) quantum dots (UbiQD, Los Alamos, USA) with zinc oleate ligands (zinc complex with oleic acid). These quantum dots are commercially available in four colours in the visible range with peak fluorescence (±10 nm) at 550 nm (green), 590 nm (yellow), 620 nm (orange), and 650 nm (red). We dissolved quantum dots in hexane to make a dispensable quantum-dot solution. The concentration and volume of quantum-dot solutions applied to anthers were tailored to suit pollen and anthers of each plant species tested (see below). Quantum-dot solutions were stored in complete darkness below 30°C inside small 2 ml clear-glass vials (9 mm thread; 12 × 32 mm, product number—29371-U; Supelco, Bellefonte, PA, USA) closed with plastic caps containing PTFE/silicone septa (9 mm polypropylene cap; PTFE/silicone septum; product number—29319-U; Supelco, Bellefonte, PA, USA). The vial septum composition is important for safe long-term storage of quantum-dot solutions as non-PTFE/silicone septa are eroded by hexane fumes, as are plain plastic caps.

### Quantum dot application to pollen

We applied quantum dots directly to individual dehisced anthers using a micropipette (0.1–2.0 µl; product code—p3942-2; Biopette, Labnet International, Edison, NJ, USA) and extra-long 10 µl pipette tips (SuperSlik(tm) 10 µL Extra Long pipet tips; product code 1165-800; Labcon, Petaluma, CA, USA). These extra-long pipette tips have a very narrow inner diameter which prevents volatile hexane from flowing out of the tip before the quantum-dot solution can be applied to pollen. When applying quantum-dot solution to anthers, we were careful to avoid direct contact between pollen and the pipette tip. We held the pipette tip as close as possible to the edge of an anther and ejected the quantum-dot solution slowly onto the anther, allowing it to gently flow across the anther and cover all pollen grains. Hexane is highly volatile (boiling point: 68°C), and therefore evaporates seconds after application, leaving behind quantum dots which putatively bind to the pollenkitt of pollen grains through lipophilic ligands.

### Reading quantum dot labels

To detect potential quantum dot labels on pollen grains requires quantum dot excitation and visualisation of the emitted quantum-dot colour. Most commercially available fluorescence microscopes are designed to visualise older generation fluorophores (e.g., green fluorescent protein, DAPI) and require modification to excitation and emission optical filters to allow visualisation of quantum dots. Moreover, most fluorescence microscopes are designed for viewing samples prepared on microscope slides. To view quantum-dot-labelled pollen grains on insects would require an additional fluorescence dissection microscope. Most pollination labs do not have fluorescence dissection microscopes at their disposal, and these microscopes are often prohibitively expensive.

However, most pollination labs already own a basic dissection microscope. We therefore designed a quantum-dot excitation box which can be placed under any standard dissection microscope allowing cost-effective visualisation of quantum dots. The box contains four LED lights (Intelligent LED Solutions, C3535 1 Powerstar Series UV LED, 390 nm, 400 mW, 125° light angle, 4-Pin) as the UV excitation source. Quantum dots are viewed through a viewport containing a UV blocking, long-pass filter (blocking wavelengths < 435 nm; Schott GG435, 50 x 50 mm) which is aligned underneath the microscope objective. The box can hold microscope slides on a drawer which slides in and out of the box (Fig. 1). To view insects, the drawer is removed and a rod with a clamp is used to hold insects in place.

**Figure 1.**
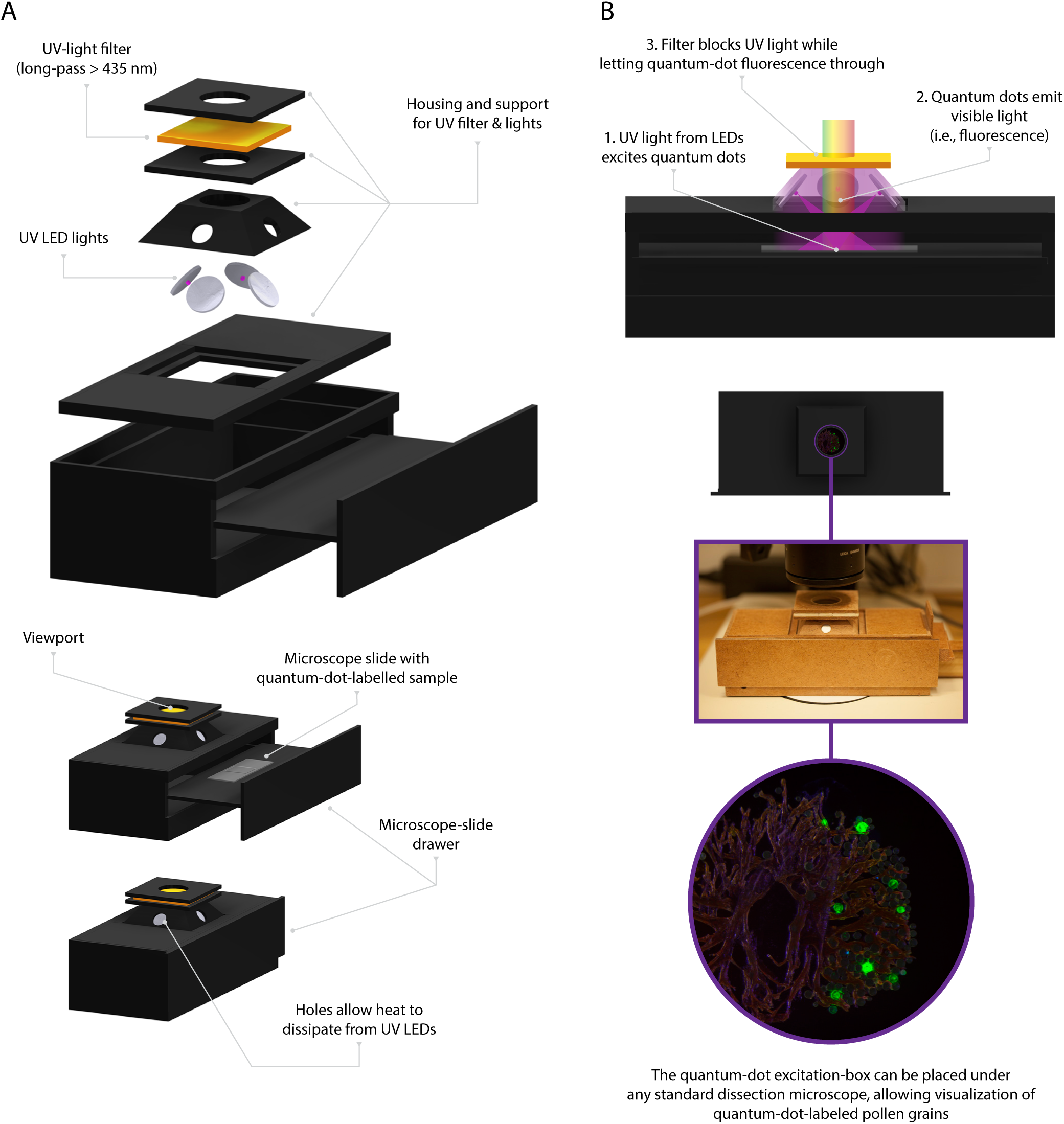
Diagram showing the components (A) of the quantum-dot excitation box and how it functions (B).

## METHOD VALIDATION

For quantum dots to be useful as pollen grain labels, they need to satisfy three key criteria: (1) quantum dots must attach directly to pollen grains; (2) the application of quantum dots to an anther should result in the labelling of most pollen grains in the anther; (3) quantum dot labels should not affect pollen grain transport.

We evaluated criteria 1 and 2 using four different plant species from four different families: *Wachendorfia paniculata* (Haemodoraceae), *Sparaxis villosa* (Iridaceae), *Arctotheca calendula* (Asteraceae), *Oxalis purpurea* (Oxalidaceae). Flowers were, collected from urban parks and gardens in Stellenbosch, Western Cape, South Africa. These four species were selected as representative of typical plants to which the quantum dot labelling technique may be applied. For each species, we determined the appropriate volume for quantum-dot application: we started by applying a 0.1 µl dose of pure hexane to an individual anther and checked whether the volume was sufficient to cover the entire anther. If the volume was too small, we increased the dosage incrementally by 0.05 µl until a single dose was sufficient to cover, but not flood, the entire anther surface. Suitable dosage volumes for the four different species were as follows: 0.30 µl per anther for *W. paniculata*; to 0.50 µl per anther for *S. villosa*; 0.35 µl per anther for *A. calendula*; and 0.15 µl per anther for *O. purpurea*. This roughly equated to the volume of anther (measured as product of the length, height, and width of the anther to the closest 0.5 mm) divided by five.

Next, we determined the appropriate quantum-dot concentration for quantum-dot application to pollen. We started by applying a 2 mg/ml (quantum-dot/hexane) solution at the ideal volume determined for each species. We then placed labelled pollen next to unlabelled pollen (separated by 3 mm) using two separate, sterile pipette tips. We examined the grains using the quantum-dot excitation-box to check whether labelled grains were easy to distinguish from unlabelled grains. Although labelled grains were distinguishable from unlabelled grains at 2 mg/ml (quantum-dot/hexane) solution, we increased the concentration of the quantum-dot solution to 5 mg/ml for all species to ensure that labelled grains were clearly distinct from unlabelled grains.

### Criteria 1: Do quantum dots attach to pollen grains?

We hypothesised that lipophilic ligands on quantum dots would attach to pollenkitt surrounding pollen grains. In the dosage trails above, it was clear that quantum dots coated pollen grains. However, this does not necessarily mean that quantum dots attached to pollen grains. To test whether quantum dots physically attach to pollen grains, we applied quantum-dot solution to five different anthers for each of the four species. We then removed the anthers and placed each inside a small centrifuge tube (0.3 ml) containing 100 µl of 70% ethanol–distilled water solution. To dislodge pollen from anthers into the ethanol solution, we vortexed each tube for two minutes. The 70% ethanol solution acts as a polar solvent and will therefore not remove pollenkitt, which is primarily hydrophobic (Pacini and Hesse, 2005), from pollen grains. If dots are physically attached to pollenkitt on pollen grains, they should remain on pollen grains when agitated in ethanol. However, if quantum dots are not physically attached to pollenkitt on pollen grains, they would be removed from pollen grains during agitation in ethanol and precipitate. After vortexing, we waited two minutes to allow potentially unattached quantum dots to precipitate while pollen grains remained suspended within the solution. We then took five 10 µl subsamples of the pollen-ethanol suspension using a micropipette and expelled each subsample onto a separate microscope slide to see if quantum dots remained attached to pollen grains using the quantum-dot excitation box.

### Criteria 2: What proportion of grains in an anther are labelled?

In addition to confirming quantum dot attachment in the experiment above, we also determined the proportion of labelled to unlabelled grains by counting every labelled and unlabelled pollen grain in each 10 µl subsample. While not all pollen grains present within anthers were counted, we assumed the random sampling of pollen grains after vortexing was representative of all pollen grains within an anther.

### Criteria 3: Do quantum dots influence pollen transport?

If quantum dots successfully attach to pollen grains, the presence of quantum dots on pollen grains may still influence how pollen grains get transported, limiting their utility as pollen-grain labels which provide biologically realistic estimates of pollen movement. To test whether quantum-dot labels influence pollen transport, we conducted multiple labelled and unlabelled pollen transfer trails for *S. villosa* using honeybees *Apis melli-fara capensis* as pollen vectors.

To compare pollen transfer dynamics of labelled and un-labelled pollen under natural conditions is unfeasible because pollen-vectors carry pollen on their bodies from previous flowers, which makes it impossible to distinguish pollen transferred from target donors and pollen from previous donors. Therefore, any comparison of labelled and unlabelled pollen transfer requires vectors to be completely clean of any pollen, which is most effectively achieved by rearing vectors in captivity.

#### Honeybee maintenance and training

We obtained ca. 400 newly-emerged adult honeybees from three semi-wild brood frames placed inside an incubator at 36°C for 48 hours. All adult honeybees were removed from brood frames before incubation to ensure that all adult honeybees taken from frames after incubation were newly-emerged. We then placed the honeybees inside a polystyrene mini-nucleus hive (Apidea: Bruck-enstrasse 6 CH-3005, Bern, Switzerland) containing pre-formed wax comb with bee bread and a constant supply of 50% sugar solution presented inside 0.3 ml centrifuge tubes. We kept the mini-nucleus hive inside a flight cage (70 × 70 × 140 cm) with a central partition that divided the cage into two equal halves. One half of the flight cage housed the bees for training and maintenance, while the other half was reserved for pollen transfer experiments (see below). The flight cage was kept indoors at 25-30°C and a 12:12 hour, light:dark cycle.

Once bees were actively flying within the flight cage (one week), we trained them to collect nectar from *S. villosa* flowers. To train bees, we placed six emasculated *S. villosa* flowers inside the flight cage for at least four hours per day. Each flower was securely attached to the top of 30cm long bamboo skewers secured to the cage floor. Flower stems were held inside small centrifuge tubes (0.3 ml) containing water. We supplemented nectar (20% w/w sucrose/tap water added to flowers in 5 µl doses) in flowers when empty to ensure that flowers remained rewarding irrespective of honeybee foraging rate. Honeybees started foraged consistently from flowers after three days of training, at which point we commenced pollen transfer experiments. Training continued throughout experiments.

#### Pollen transfer experiments

At the start of each experimental day, we picked unopened *S. villosa* flowers in the morning and randomly split them into two groups, donors and recipients, in a 1:10 ratio. All recipients were emasculated in the morning prior to anther dehiscence and used on the following day when stigmas were mature and receptive. In addition, we checked the stigmas of all recipients for any pollen grains under a dissection microscope. All flowers with stigmatic pollen were discarded. We removed the stigmas of donor flowers in the morning prior to anther dehiscence and assigned them randomly to one of two treatments: labelled or unlabelled pollen. Once anthers were fully dehisced, we either left the flowers as they were (unlabelled pollen) or applied quantum-dot solution to the anthers as described before (labelled pollen).

For each pollen transfer trial, we placed 11 *S. villosa* flowers in a line perpendicular to the cage partition (spaced 5 cm apart) on the experimental side of the flight cage. The first flower in the line acted as the pollen donor, while the next 10 flowers behind acted as pollen recipients. The donor flower was placed 2 cm away from a small door (5 x 10 cm) in the cage partition. This door could be opened and closed from the outside of the cage, to allow or prevent bees passing from one part of the cage to the other. Flowers were attached to bamboo skewers as before, but we allowed part of the bamboo skewer to extend above flowers which enabled us to cover individual flowers with a small plastic cup and prevent bees from visiting any flower more than once.

To start a pollen transfer trial, we opened the door in front of the donor flower. At the same time, we held a piece of cardboard just behind the donor flower so that honeybees could not see the recipient flowers. We waited for a honeybee to fly through the door and visit the donor flower, after which we closed the door and removed the cardboard blocking the recipient flowers. Once the bee finished visiting the donor flower, we covered it with a plastic cup to prevent repeat visitation. Thereafter, we ensured that all flowers received only one visit by covering flowers with a plastic cup immediately after being visited. The plastic cups were pre-numbered allowing us to determine the visitation sequence afterwards. Once the honeybee finished visiting all of the flowers, we captured and killed it to ensure that pollen was not transported back to the training area. We completed 15 pollen transfer trials for each treatment. Only three trials out of 30 resulted in visits to 9 recipients instead of all 10 (unlabelled: 2; labelled: 1).

After a trial was complete, we recorded the position in the transfer sequence for each flower and the treatment applied to the donor flower. We also measured the closest distance between the stigma and the ventral corolla surface (stigma height), as this may influence the likelihood of pollen being transferred. Additionally we measured the closest distance between the donor anthers and the ventral corolla surface (anther height) as this may influence the likelihood of pollen being picked up by the honeybee. We harvested the stigmas of each recipient and placed them on individual microscope slides. To prepare stigma slides, we squashed unlabelled stigmas under a cover slip with melted Fuschin gel, while labelled stigmas were squashed under a cover slip without a mounting medium and the edges of the cover slip secured and sealed using transparent sticky tape. Stigma slides were stored in the freezer at −20°C until pollen could be counted. We counted pollen using a standard dissection microscope with labelled pollen visualised inside the quantum-dot excitation box.

We modelled pollen transfer using a non-linear regression of the number of pollen grains deposited on stigmas (pollen count) as an exponential decay function of visit-sequence number (*visit seq*.), stigma height (*stigma h*.), and anther height (*anther h*.):

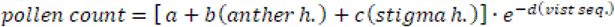

The expression within the square brackets determines the number of pollen grains at the first visit in the sequence, while the expression to the right of the square brackets determines the rate of decay (pollen depletion) as a function of visit sequence number. Therefore *a*, represents an independent intercept, and *b* and *c* are expected to be negative because increased stigma and anther heights may result in decreased transfer of pollen due to poor contact with honeybees. The coefficient *d* is expected to be positive and controls the magnitude of pollen-transfer decay with increasing visit sequence number.

To determine if treatment had an effect on pollen carry-over, we first ran two non-linear regressions using the model structure above: in the first model, the coefficients were determined separately for each treatment (i.e., the model assumed treatment had an effect on pollen transfer); the second model ignored treatment and determined the coefficients across treatments. We then performed a Chi-square goodness-of-fit test comparing the residual sum of squares for the two models, to determine whether fitting different coefficients based on treatment resulted in a significant reduction in error compared to the model which ignored treatment effects. Models were computed in R using the “nls” function (R Core Team, 2017).

## RESULTS & DISCUSSION

### Criteria 1: Do quantum dots attach to pollen grains?

Quantum dots remained attached to pollen grains even after agitation in 70% ethanol, in all subsamples, for all four species (n = 25 subsamples per species). This strongly suggests that quantum dots attached to pollen grains through a lipophilic interaction between their oleic acid ligands and the lipid-rich pollenkitt surrounding pollen grains. Nearly all animal-pollinated angiosperms have pollen grains surrounded by pollenkitt (Pacini and Hesse, 2005). Although, pollenkitt composition varies, lipids remain the primary constituent of pollenkitt. Therefore, quantum dots will likely bind to pollen grains of most animal-pollinated species. Another sticky pollen-coat substance called tryphine is found only in Brassicaceae (Pacini and Hesse, 2005). While it is functionally similar to pollenkitt, it is composed of both lipophilic and hydrophilic substances (Dickinson and Lewis, 1973). The suitability of quantum dots as pollen labels in species with tryphine pollen coatings remains unconfirmed. However, the wide array of ligands available for attachment to quantum dots will allow tryphine-specific ligands to be developed if oleic acid ligands are not suitable.

### Criteria 2: What proportion of grains in an anther are labelled?

The majority of pollen grains in each subsample were labelled by quantum dots. No unlabelled grains were found in any of the subsamples for *A. calendula*, while nearly all grains were labelled for *W. paniculata* (mean ± SE: 0.97±0.01) and *S. villosa* (0.92 ± 0.01) (Fig. 2). The proportion of grains labelled for *O. purpurea* was comparatively low, but most grains were still labelled (0.74 ± 0.02). Anthers of *O. purpurea* release pollen through very narrow apertures and do not fully open upon dehiscence. This semi-closed anther structure may explain why a smaller proportion of *O. purpurea* grains were labelled relative to the other species. It may therefore be difficult to label all pollen grains in species with closed anther structures (e.g., buzz-pollinated anthers). However, for most applications, labelling all pollen grains in an anther may be unnecessary, since comparisons of pollen transfer will likely be relative. For studies comparing pollen transfer across species, we recommend quantifying the proportion of pollen grains labelled in anthers following methods in this paper and applying these proportions as species-specific correction factors for quantitative assessments of pollen movement.

**Figure 2.**
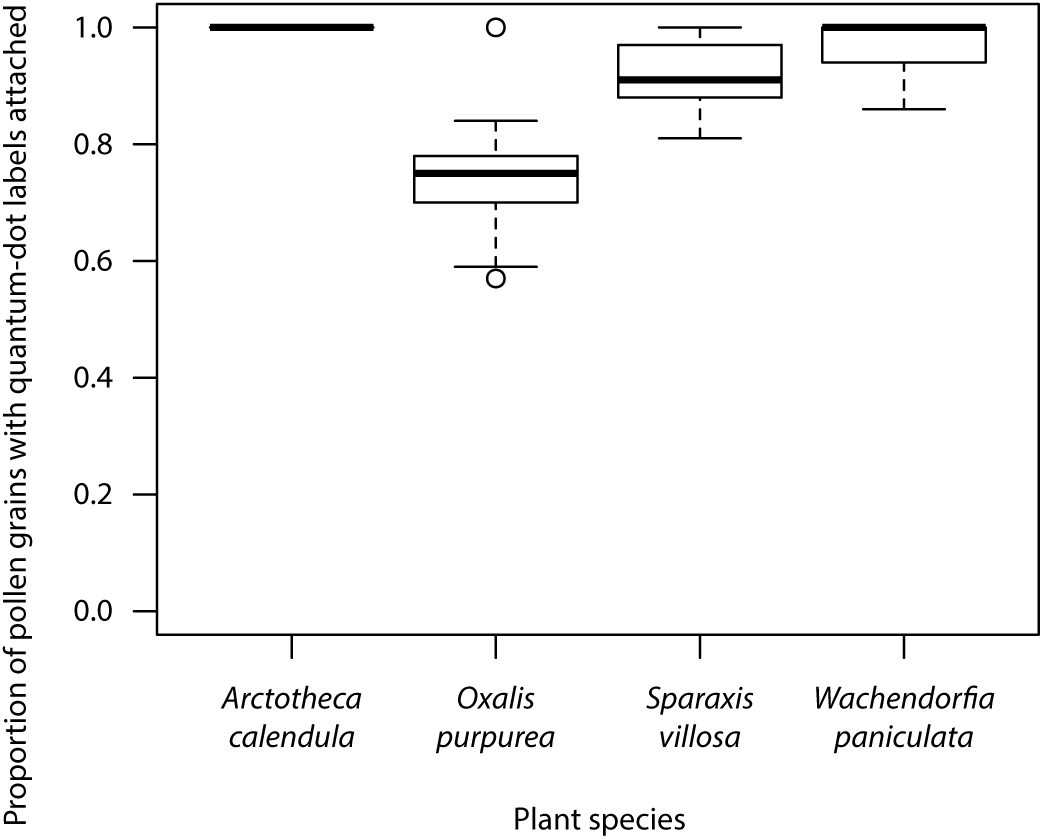
Boxplot of the proportion of pollen grains labelled with quantum dots after agitation in 70% ethanol.

### Criteria 3: Do quantum dots influence pollen transport?

Differentiating between unlabelled and labelled treatments did not significantly decrease model error (F = 0.8705; p = 0.482). Stigma height (*c*) and visit sequence number (*d*) were the main determinants of pollen transfer (*c*: mean ± SE = −92.38 ± 8.30, t = −11.13, p < 0.0001; *d*: mean ± SE = 0.27 ± 0.02, t = 12.62, p < 0.0001). These results suggest that using quantum dots as pollen labels did not affect pollen carryover in *S. villosa*. This may be a consequence of the small amount of quantum dots required to label almost all pollen grains in anthers. For *S. villosa*, only 2.5 µg of quantum dots were applied to an entire anther. The mass attached to pollen grains is likely even less than that (some quantum dots remain on anthers and in the pipette tip). Although no attempt was made to estimate the mass of quantum-dot labels attached to an individual pollen grain, it is likely a tiny fraction of the pollen grain’s total mass and does not appear to influence pollen transfer between flowers (Fig. 3).

**Figure 3.**
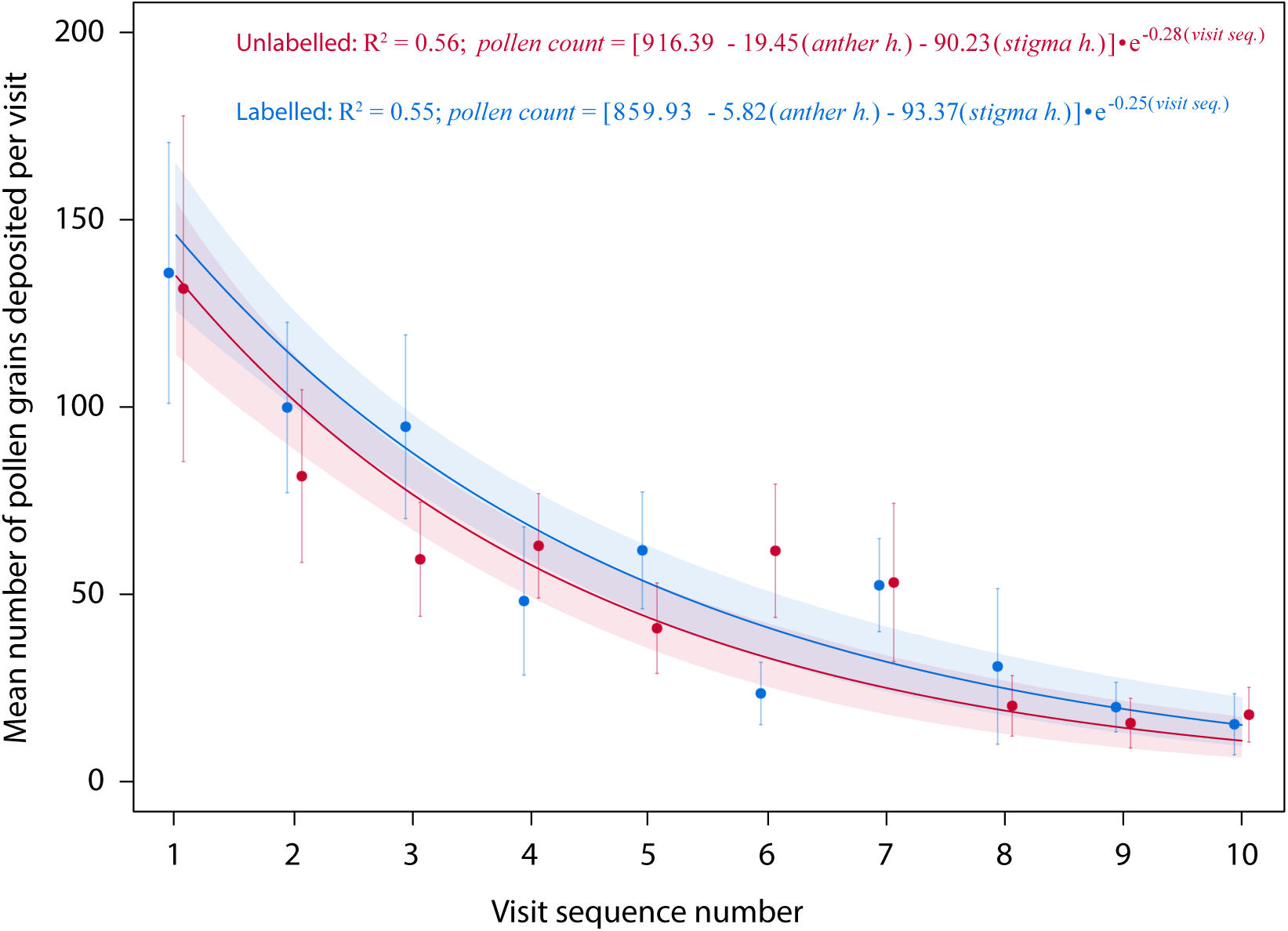
The mean number of pollen grains deposited per visit as a function of visit sequence number. Dots and errors bars (red: unlabelled pollen; blue: labelled pollen) indicate mean and standard error. The red (unlabelled pollen) and blue (labelled pollen) lines indicate the predicted mean pollen deposition from non-linear regressions, after controlling for stigma and anther height. Shaded areas show the 95% confidence intervals for model estimates. The equations for both regressions are shown with mean coefficient estimates for equation 1.

### Limitations and future improvements

#### Limited colours

Currently, the number of distinguishable quantum dot colours that are commercially available is limited to four. This is likely not a serious limitation for most pollen movement studies; however, having only four colours may limit simultaneous assessments of pollen movement from more than four individual plants in close proximity. The number of quantum dot colours may increase with time since techniques for producing non-toxic quantum dots are being developed at a rapid pace, and more long-wavelength colours (blue and violet) may soon be available for non-toxic quantum dots.

It may also be possible to create unique pollen labels by combining different quantum dot colours. The use of multi-coloured quantum-dot-beads as individually attached multiplexed labels have been demonstrated for biomolecules (Han *et al*., 2001). However, binding individual beads (1-2 µm) to individual pollen grains would require the development of species-specific binding protocols which negates the universality of the current method. A simpler method to achieve multiplexed quantum-dot pollen labels may be to mix two or three quantum-dot colours together to make up a colour code. For example, an equal mix of red and green quantum dots would have the code RG. When viewed normally through a quantum-dot excitation-box, the visible colour of pollen grains labelled with the RG mixture, would simply appear yellow. However, the RG code can be read by adding band-pass filters to the view-port of the microscope. If a green band-pass filter is applied, true yellow-labelled pollen grains will become invisible, while RG-labelled grains would appear green. Similarly, by applying a red band-pass filter, the red component of RG-labelled grains will become visible. With four quantum-dot colours and four band-pass filters, 14 unique quantum-dot-labels could be created from single, bi-colour, and tri-colour codes combined. Preliminary trails suggest that multiplexed quantum-dot labels are possible (unpublished data), but the efficacy of multiplexed colour codes will depend on the emission bandwidth of quantum dots and the cut-off slope of the bandpass filters. Individual quantum dots have extremely narrow emission spectra, but a mix of quantum dots will have a wider emission spectrum due to variation in size of quantum dots within the mixture, which may vary depending on the precision of size-control that quantum-dot producers exercise. If the emission spectrum of different quantum dot colours overlaps too much, the multiplexing method will have limited success. Similarly, poor quality band-pass filters may not isolate specific colours well enough, leading to false confirmations of colours.

Although multiplexed quantum-dot codes may increase the number of unique pollen labels, the effort required to read these codes will be substantial. Because reading codes will require switching between optical filters and keeping track of the positions and colours of various individual pollen grains, this method is only feasible if photographs are taken of samples through each band-pass filter and codes assigned to individual pollen grains afterwards.

## CONCLUSION

Here we have demonstrated a method which promises to arm pollination biologists with the ability to rapidly label and subsequently track the fates of pollen grains in the field for perhaps the majority of angiosperm species. For more than 150 years we have been largely unable to address various aspects of floral evolution because of the lack of techniques available to study pollen movement. These include, but are not limited to: the magnitude and frequency of pollen loss during various stages of the pollen export process (Inouye *et al*., 1994); the importance of vector-mediated pollen-movement isolation as a speciation and diversification mechanism in angiosperms (Armbruster, 2014); the functional effects of heterostyly and enantiostyly on pollen movement between floral morphs (Barrett and Shore, 2008); and, the structure and competitive implications of the various pollen landscapes that form on vectors as a result of sequential visits to competing conspecifics and heterospecifics (Harder and Wilson, 1998; Muchhala *et al*., 2010). The application of quantum-dot nanotechnology for pollen labelling may finally allow direct assessments of pollen movement in most angiosperms, adding to our understanding of a neglected but vital aspect of floral evolution.

